# ACE2 coding variants in different populations and their potential impact on SARS-CoV-2 binding affinity

**DOI:** 10.1101/2020.05.08.084384

**Authors:** Fedaa Ali, Menattallah Elserafy, Mohamed H. Alkordi, Muhamed Amin

## Abstract

The susceptibility of different populations to the SARS-CoV-2 infection is not yet understood. A deeper analysis of the genomes of individuals from different populations might explain their risk for infection. In this study, a combined analysis of ACE2 coding variants in different populations and computational chemistry calculations are conducted in order to probe the potential effects of ACE2 coding variants on SARS-CoV-2/ACE2 binding affinity. Our study reveals novel interaction data on the variants and SARS-CoV-2. We could show that ACE2-K26R; which is more frequent in the Ashkenazi Jewish population decrease the electrostatic attraction between SARS-CoV-2 and ACE2. On the contrary, ACE2-I468V, R219C, K341R, D206G, G211R were found to increase the electrostatic attraction and increase the binding to SARS-CoV-2; ordered by the strength of binding from weakest to strongest. I468V, R219C, K341R, D206G and G211R were more frequent in East Asian, South Asian, African and African American, European and European and South Asian populations, respectively. SARS-CoV-2/ACE2 interface in the WT protein and corresponding variants is showed to be a dominated by van der Waals (vdW) interactions. All the mutations except K341R induce an increase in the vdW attractions between the ACE2 and the SARS-CoV-2. The largest increase of is observed for the R219C mutant.

## Introduction

Despite numerous reports on the clinical manifestation of the recent novel coronavirus disease 2019 (COVID-19), including pathogenicity, diagnosis, and recommended treatment regimes, understanding of the mechanisms of infection, the virus-host interactions, and transmission is still in its early stage^1–4^. The risk of certain populations for the infection is also still not well understood and remains under investigation. After the identification of the angiotensin-converting enzyme 2 (ACE2) as the SARS-CoV-2 receptor^5^, a large number of reports have emerged trying to identify the pathology of the disease, and thus provide guidance to the efforts to design small molecules inhibitors^3,5^. Some recent studies attempted to probe the comorbidity among infected patients, specifically patients with hypertension, diabetes mellitus, coronary heart diseases, and cerebrovascular disease that showed increased risk. Whether ACE2 polymorphisms linked to the aforementioned diseases or administration of ACE2 modulating drugs have larger impact on the severity of infection is still a question to be investigated^6^.

Communications between the pathogen and the host is mainly governed by the different proteinprotein electrostatic interactions^7,8^. These proteins need to bind in a specific way for proteinprotein complexes formation and for the signals to be effectively transmitted. The existence of residues that are involved in energetically favorable interactions at the protein-protein interface leads to enhanced interactions leading to viral-host cells fusion, marking the onset of the infection^5,9^. Therefore, we opted herein to investigate the nature of the interactions at the SARS-CoV-2/ACE2 interface, and how the subtle variations in the structure of ACE2, induced by certain mutations, can alter the binding of SARS-CoV-2 in different populations. This was attempted here by analyzing different ACE2 missense variants that code for ACE2-K26R, ACE2-I468V, ACE2-R219C, ACE2-K341R, ACE2-D206G, ACE2-G211R and the strength of their interaction with the spike protein (S-protein) of SARS-COV-2 using Molecular Dynamics and Monte Carlo Sampling, which are dominated by electrostatics. The electrostatics interactions are the dominant factor in protein-protein interactions^10^, where classical treatment of these interactions in solution are described by the Poisson-Boltzmann equation (PBE).

Our data demonstrates an increase in the strength of the binding of the variants; in the order mentioned, with K26R showing the least favorable binding and G211R showing the most favorable binding. We analyzed the frequencies of the alleles in different populations to identify the population in which each variant is more frequent. Our data help in identifying variations of appreciable frequencies and the implications for each in affecting the strength of the virus-receptor interactions in individuals of different populations.

## Materials and methods

Classical electrostatic calculations based on solving Poisson-Boltzmann (PB) equation is used to investigate the interaction energies between SARS-CoV-2 and ACE2 for different mutated ACE2 structures. All continuum electrostatics calculations were performed by the freely available software “Multi-Conformer Continuum electrostatics (MCCE)”^11^. Where that interaction energies are modeled by treating each residue as separate fragments with integer charges, which are interacting with each other by means of electrostatic and Lennard-Jones potentials^11^. The Poisson-Boltzmann (PB) equation is numerically solved by DELPHI software, which is integrated within the MCCE code, to compute self-energies of each conformer and pairwise interaction energies between conformers and between conformers and backbone. Upon energy calculations, the energy look-up table is built, and theoretical titration is modeled by obtaining Boltzmann distribution of microstates, using the Monte-Carlo approach. Where that microstates of the structures are defined depending on rotamers and protonation states of the different amino acids. Where that the protein interior are considered as highly polar media with low dielectric constant (*ε_prot_* = 4), whereas the solvent (water) is treated as a continuum medium with high dielectric constant (*ε_wat_* = 80).

Initial coordinates are defined according to the X-ray crystal structure of the wild type (WT) human ACE2 with the receptor-binding domain RDB of the S protein of SARS-CoV-2 (PDB ID: 6M17) from the protein data bank^5^. The structure of the Neutral amino acid transporter B^0^AT1 was removed from the PDB file, as it is far from the SARS-CoV-2/ACE2 binding site. Furthermore, the structure of ACE2 was trimmed by removing residues from P733 to the end of the chain. The WT residues of ACE2 at positions 26, 206, 211, 219, 341 and 468 were mutated to Arg (R), Gly (G), Arg (R), Cys (C), Arg (R) and Val (V), respectively. The sidechains in the WT structure was replaced with the sidechains of the mutants using MCCE (Multi Conformer Continuum Electrostatics) (Fig. 1). A set of different conformers of the sidechains were generated for WT and mutated structures by rotating the single bonds in the sidechains by 60^0^ degrees. In addition, conformers were created for charged amino acids according to their possible protonation states. Then, these conformers were subjected to Monte Carlo sampling to obtain the Boltzmann distribution based on the electrostatic interactions, which are calculated using DELPHI. The most occupied conformers in the generated Boltzmann distributions were selected and subjected to molecular dynamics optimization using openMM. Finally, the interaction energies were computed for the optimized structures using MCCE. The electrostatics and van der Waals interactions between aminoacids in the RBD of SARS-CoV-2 and ACE2 were analyzed using python-based code.

**Figure 1.**
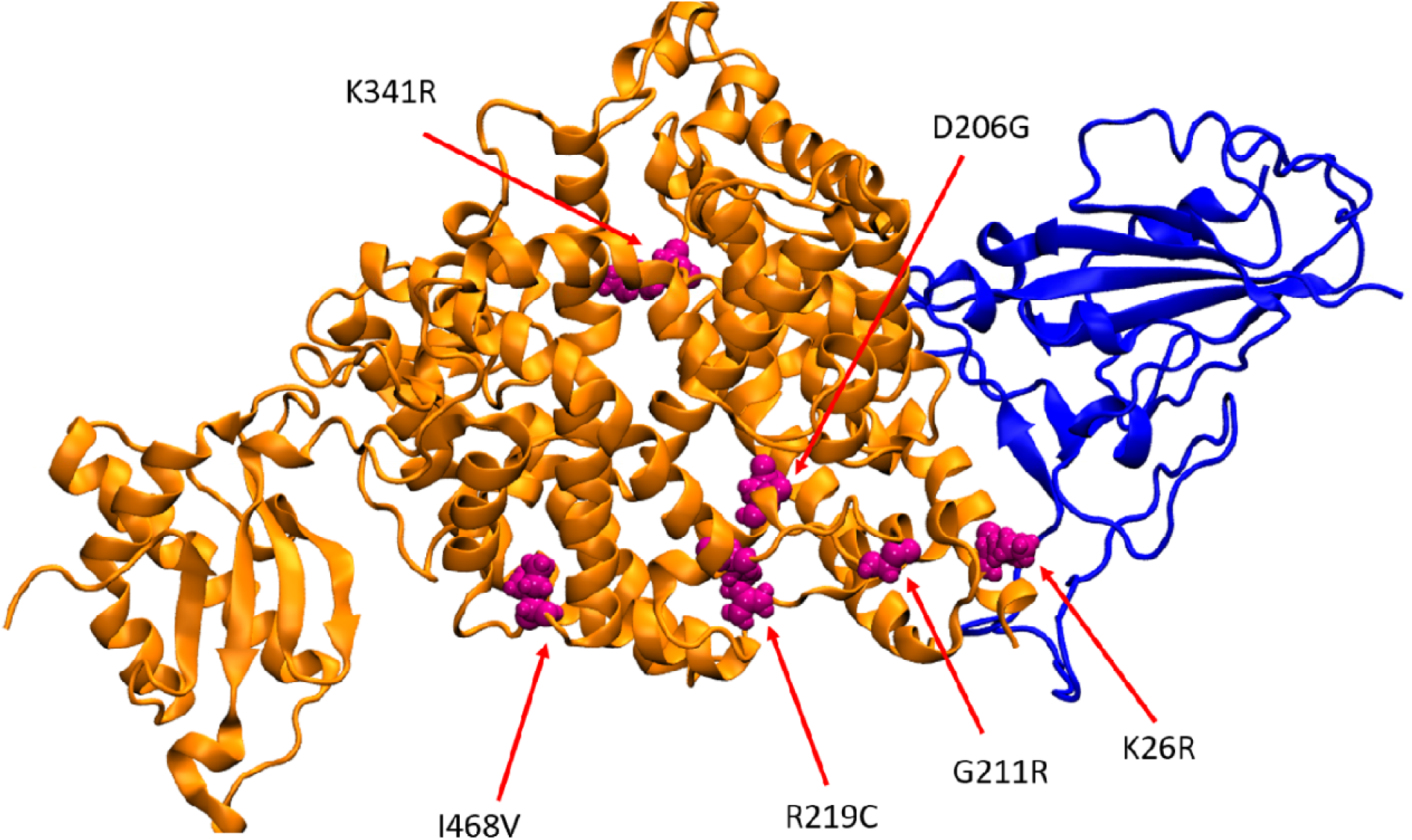
The secondary structure of the ACE2 protein (shown in orange) and the RBD of the SARS-CoV-2 (shown in blue). The mutated amino acids are shown in magenta spheres (6M17)^5^.

## Results and discussion

In an attempt to better understand the susceptibility of different populations to infection by SARS-CoV-2, we gathered data on ACE2 missense variants from different projects and databases that aggregate allele frequencies (AF). These projects and databases included Single Nucleotide Polymorphism Database (dbSNP)^12,13^, 1000 genomes project phase 3 (1KGP3)^14^, Allele Frequency Aggregator (ALFA project)^a^, Exome Aggregation Consortium (ExAC)^15^, genome Aggregation Database (gnomAD)^b,16^, GO exome sequencing project (ESP)^c^, Trans-Omics for Precision Medicine (TopMed)^17^, in addition to a recent study reporting data from the China Metabolic Analytics Project (ChinaMAP, under reviewing) and other populations^18^.

SARS-CoV-2 was reported to bind to human ACE2 via different ACE2 residues; Q24, D30, H34, Y41, Q42, M82, K353 and R357^5^. We analyzed all ACE2 coding variants reported and selected the variants that are close to the interaction site to test for changes in their binding interactions to SARS-CoV-2. To focus on the most frequent variants, we did not include variants of very low allele count in all populations. The variants we tested were K26R, D206G, G211R, R219C, K341R and I468V. Table S1 shows the frequency of each allele in the African, African American, American, Ashkenazi Jewish, Asian (All, ChinaMAP, East Asian, Han Chinese South, Han Chinese Beijing and South Asian), European (All, European American, Finnish and non-Finnish) and Latino.

### Variations in ACE2 affect its interaction with SARS-CoV-2

The electrostatic and the van der Waal contribution to the interaction energies of SARS-CoV-2/ACE2 were compared between single mutated and WT protein at pH =7 (Table 1). In contrary to K26R, most of the mutated structures exhibit a negative shift in the total interaction energy compared to the WT structure (Native) i.e. more electrostatic attraction with the spike protein of SARS-CoV-2. Based on our calculations, G211R mutant is shown to induce the largest increase in the binding energy between SARS-CoV-2 and ACE2, where the binding is more favorable by ∼ 7.6 Kcal/mol than the WT. However, the K26R mutant causes a decrease in the binding energies by ∼ 2.1 Kcal/mol (Table 1).

**Table 1.** The interaction energies between the SARS-CoV-2 and ACE2

| **Table 1.** The interaction energies between the SARS-CoV-2 and ACE2 |  |  |  |  |
| --- | --- | --- | --- | --- |
|  | Electrostatics<br>(Kcal/mol) | vdW<br>(Kcal/mol) | Total<br>(Kcal/mol) | Population |
| G211R | -12.4 | -35.4 | -47.8 | European<br>South Asian |
| D206G | -8.8 | -36.1 | -44.9 | European |
| K341R | -14.2 | -29.3 | -43.4 | African/African<br>American |
| R219C | -4.6 | -38.3 | -42.9 | South Asian |
| I468V | -4.7 | -37.4 | -42.1 | East Asian |
| **Native** | **-9.6** | **-30.1** | **-40.2** |  |
| K26R | -3.7 | -34.4 | -38.1 | Ashkenazi<br>Jewish |
| The entries are sorted from lowest to the highest interaction energies. |  |  |  |  |

The substitution of Lys (K) to Arg (R) at positions 211 and 341 was shown to increase the electrostatic attraction between SARS-CoV-2 and ACE2 by 1.29-fold and 1.47-fold in comparison to the WT, respectively. Oppositely, substitution of R at position 26 was shown to reduce the electrostatic attraction between SARS-CoV-2 and ACE2 to 2.59 Kcal/mol lower than that in WT. In addition, the D206G mutant showed similar electrostatic interactions to the WT. However, the R219C, I468V mutants showed significant decrease in the electrostatic attraction to by ∼5 Kcal/mol compared to the WT (Table 1). SARS-CoV-2/ACE2 interface in the WT protein and corresponding mutants is showed to be a dominated by vdW interactions, which accounts for more than 60% of the interaction energy. All the mutations except K341R induced an increase in the vdW attractions between the ACE2 and the SARS-CoV-2. The largest increase of ∼ 8 Kcal/mol is observed for the R219C mutant (Table 1).

Our results, shown in Fig. 2, demonstrate large variations in calculated binding energy in different ACE2 variants (identified in the table 1) towards SARS-CoV-2 S protein. Although the calculated variations in binding energies are appreciable, as compared to the control, we hypothesize that even within the same population mutations in ACE2 in certain individuals can similarly dramatically influence the binding site configuration and ACE2/SARS-CoV-2 binding affinity.

**Figure 2.**
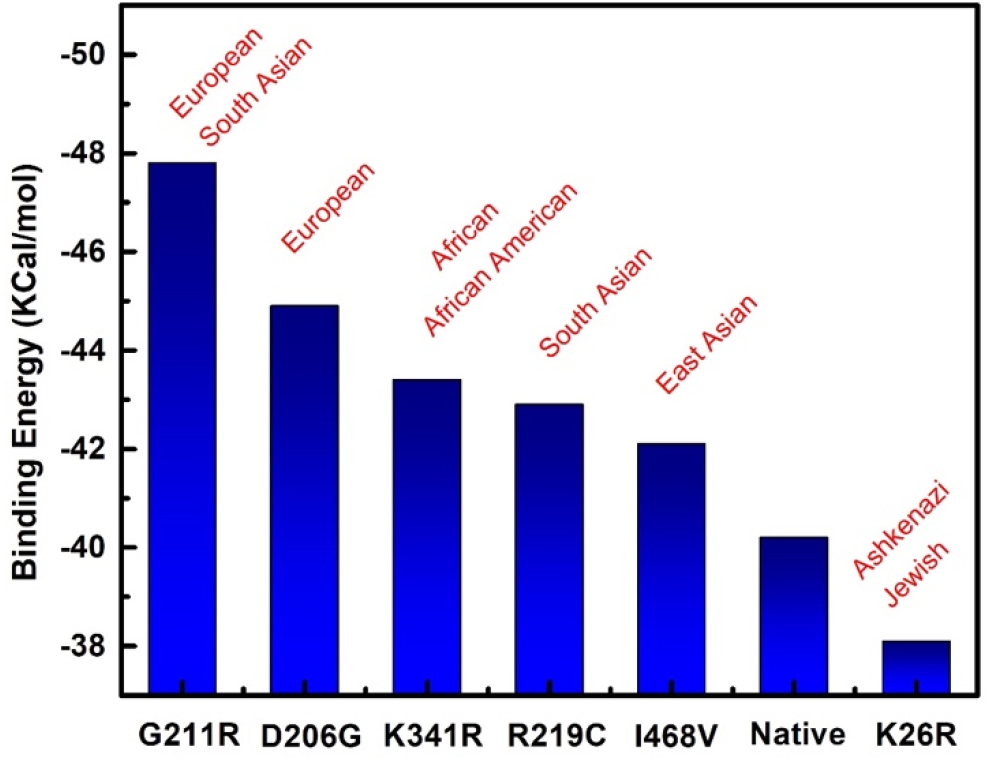
Representation of Binding energies calculated in Table 1 and the corresponding population with the highest allele frequency.

### How ACE2 variants can impact the susceptibility of different population to SARS-CoV-2 infection?

The K26R variant which decreases the ACE2-virus binding was found to be most frequent in the Ashkenazi Jewish population (1.2%). On the contrary, the Asian population had the lowest allele frequency for the single nucleotide variant (SNV) coding for K26R. In addition, the frequency of I468V which enhance ACE2 interaction with SARS-CoV-2 was observed in more than 1% of the East Asian population. Interestingly, I468V was not detected in the African, African American, American, Ashkenazi Jewish and Latino populations and was very rare in the European population. R219C was most frequent in the south Asian population (0.1%), but was not detected in other Asian subpopulations and was much less frequent or lacking in other populations. The K341R, which enhance the virus-receptor binding was most frequent in the African and African American populations; the average of the AF reported for the populations was ∼ 0.4% and ∼ 0.5%, respectively. Finally, the D206G and G211R, which showed the strongest interaction with ACE2 were most abundant in the European population, mounting to 0.06% and ∼ 0.2%, respectively. The G211R was also abundant in the South Asian population ∼ 0.2%.

Our findings give indications regarding the populations that could be more prone to SARS-CoV-2 infection due to enhanced binding affinity. Nevertheless, the need for proper infection, death and recovery statistics is needed to reach a definite conclusion. K26R and I468V are considered of intermediate frequencies (>1%) in the Ashkenazi Jewish and East Asian population, respectively. Therefore, we would refer to them as Single Nucleotide Polymorphism (SNPs) in the respective populations. However, the other variants are considered of rare frequencies (<1%)^19,20^. Nevertheless, in the era of Individualized Genomics, the individual differences in the same population should also be taken into account as rare variants might exist affecting the virus binding to ACE2 in many individuals.

The possibility of having individuals baring more than one of the mutations that enhance the binding is also still an open question, that will be answered via sequencing of ACE2 in more individuals of the aforementioned populations. The presence of variants for the serine protease TMPRSS2; required for S protein priming, in different populations could also account for differences in viral entry and pathogenesis^9^.

## Conclusion

Our data suggests that certain populations might be more affected by SARS-CoV-2; depending on the frequency of the respective variants. To better understand the susceptibility of individuals from different populations to SARS-CoV-2 and their risk of infection, large scale sequencing projects should be performed. Integrating population genetics will provide a new insight into the mystery of susceptibility, infection, pathogenicity and mortality in different regions. Sequencing of ACE2 in patients with the most severe conditions in every population will also provide a more reliable conclusion. The sequencing projects will not only help us test the association between ACE2 variants and risk/severity of infection in certain populations, it will also allow more accurate analysis of variants for other host genes involved in the viral entry and pathogenicity^21^. Finally, our findings can potentially guide future attempts to devise small molecules inhibitors, designed specifically to disrupt the interaction between the strongest ACE2 binding variants and the S protein of SARS-CoV-2^2^.

## Supporting information

Optimized MD/MC structures

## Footnotes

a L. Phan, Y. Jin, H. Zhang, W. Qiang, E. Shekhtman, D. Shao, D. Revoe, R. Villamarin, E. Ivanchenko, M. Kimura, Z. Y. Wang, L. Hao, N. Sharopova, M. Bihan, A. Sturcke, M. Lee, N. Popova, W. Wu, C. Bastiani, M. Ward, J. B. Holmes, V. Lyoshin, K. Kaur, E. Moyer, M. Feolo, and B. L. Kattman. “ALFA: Allele Frequency Aggregator.” National Center for Biotechnology Information, U.S. National Library of Medicine, 10 Mar. 2020, www.ncbi.nlm.nih.gov/snp/docs/gsr/alfa/.

b https://gnomad.broadinstitute.org/

c Exome Variant Server, NHLBI GO Exome Sequencing Project (ESP), Seattle, WA (URL: http://evs.gs.washington.edu/EVS/) [date (month, yr) accessed].

## Notes

### Competing Interest Statement

The authors have declared no competing interest.

## References

(1) Guo, Y. R.; Cao, Q. D.; Hong, Z. S.; Tan, Y. Y.; Chen, S. D.; Jin, H. J.; Tan, K. Sen; Wang, D. Y.; Yan, Y. The Origin, Transmission and Clinical Therapies on Coronavirus Disease 2019 (COVID-19) Outbreak-An Update on the Status. Mil. Med. Res. 2020, 7 (1), 1–10. https://doi.org/10.1186/s40779-020-00240-0.

(2) Li, H.; Zhou, Y.; Zhang, M.; Wang, H.; Zhao, Q.; Liu, J. Updated Approaches against SARS-CoV-2. Antimicrob. Agents Chemother. 2020, No. March. https://doi.org/10.1128/AAC.00483-20.

(3) Lin, L.; Lu, L.; Cao, W.; Li, T. Hypothesis for Potential Pathogenesis of SARS-CoV-2 Infection–a Review of Immune Changes in Patients with Viral Pneumonia. Emerging Microbes and Infections. 2020. https://doi.org/10.1080/22221751.2020.1746199.

(4) Al-Tawfiq, J. A.; Al-Homoud, A. H.; Memish, Z. A. Remdesivir as a Possible Therapeutic Option for the COVID-19. Travel Medicine and Infectious Disease. 2020. https://doi.org/10.1016/j.tmaid.2020.101615.

(5) Yan, R.; Zhang, Y.; Li, Y.; Xia, L.; Guo, Y.; Zhou, Q. Structural Basis for the Recognition of SARS-CoV-2 by Full-Length Human ACE2. Science (80-.). 2020, 367 (6485), 1444–1448. https://doi.org/10.1126/science.abb2762.

(6) Fang, L.; Karakiulakis, G.; Roth, M. Are Patients with Hypertension and Diabetes Mellitus at Increased Risk for COVID-19 Infection? Lancet Respir. Med. 2020, 8 (4), e21. https://doi.org/10.1016/S2213-2600(20)30116-8.

(7) Norel, R.; Sheinerman, F.; Petrey, D.; Honig, B. Electrostatic Contributions to Protein-Protein Interactions: Fast Energetic Filters for Docking and Their Physical Basis. Protein Sci. 2008. https://doi.org/10.1110/ps.12901.

(8) Sheinerman, F. B.; Norel, R.; Honig, B. Electrostatic Aspects of Protein-Protein Interactions. Current Opinion in Structural Biology. 2000. https://doi.org/10.1016/S0959-440X(00)00065-8.

(9) Hoffmann, M.; Kleine-Weber, H.; Schroeder, S.; Krüger, N.; Herrler, T.; Erichsen, S.; Schiergens, T. S.; Herrler, G.; Wu, N. H.; Nitsche, A.; et al. SARS-CoV-2 Cell Entry Depends on ACE2 and TMPRSS2 and Is Blocked by a Clinically Proven Protease Inhibitor. Cell 2020, 181 (2), 271–280.e8. https://doi.org/10.1016/j.cell.2020.02.052.

(10) Honig, B.; Nicholls, A. Classical Electrostatics in Biology and Chemistry. Science (80-.). 1995. https://doi.org/10.1126/science.7761829.

(11) Song, Y.; Mao, J.; Gunner, M. R. MCCE2: Improving Protein PKa Calculations with Extensive Side Chain Rotamer Sampling. J. Comput. Chem. 2009, 30 (14), 2231–2247. https://doi.org/10.1002/jcc.21222.

(12) Sherry, S. T.; Ward, M.; Sirotkin, K. DbSNP-Database for Single Nucleotide Polymorphisms and Other Classes of Minor Genetic Variation. Genome Research. 1999. https://doi.org/10.1101/gr.9.8.677.

(13) Smigielski, E. M. DbSNP: A Database of Single Nucleotide Polymorphisms. Nucleic Acids Res. 2000. https://doi.org/10.1093/nar/28.1.352.

(14) Auton, A.; Abecasis, G. R.; Altshuler, D. M.; Durbin, R. M.; Bentley, D. R.; Chakravarti, A.; Clark, A. G.; Donnelly, P.; Eichler, E. E.; Flicek, P.; et al. A Global Reference for Human Genetic Variation. Nature. 2015. https://doi.org/10.1038/nature15393.

(15) Karczewski, K. J.; Weisburd, B.; Thomas, B.; Solomonson, M.; Ruderfer, D. M.; Kavanagh, D.; Hamamsy, T.; Lek, M.; Samocha, K. E.; Cummings, B. B.; et al. The ExAC Browser: Displaying Reference Data Information from over 60 000 Exomes. Nucleic Acids Res. 2017. https://doi.org/10.1093/nar/gkw971.

(16) Karczewski, K. J.; Francioli, L. C.; Tiao, G.; Cummings, B. B.; Alföldi, J.; Wang, Q.; Collins, R. L.; Laricchia, K. M.; Ganna, A.; Birnbaum, D. P.; et al. The Mutational Constraint Spectrum Quantified from Variation in 141,456 Humans. bioRxiv 2019. https://doi.org/10.1101/531210.

(17) Taliun, D.; Harris, D. N.; Kessler, M. D.; Carlson, J.; Szpiech, Z. A.; Torres, R.; Taliun, S. A. G.; Corvelo, A.; Gogarten, S. M.; Kang, H. M.; et al. Sequencing of 53,831 Diverse Genomes from the NHLBI TOPMed Program. bioRxiv 2019. https://doi.org/10.1101/563866.

(18) Cao, Y.; Li, L.; Feng, Z.; Wan, S.; Huang, P.; Sun, X.; Wen, F.; Huang, X.; Ning, G.; Wang, W. Comparative Genetic Analysis of the Novel Coronavirus (2019-NCoV/SARS-CoV-2) Receptor ACE2 in Different Populations. Cell Discovery. 2020. https://doi.org/10.1038/s41421-020-0147-1.

(19) Agarwala, V.; Flannick, J.; Sunyaev, S.; Altshuler, D. Evaluating Empirical Bounds on Complex Disease Genetic Architecture. Nat. Genet. 2013. https://doi.org/10.1038/ng.2804.

(20) Bomba, L.; Walter, K.; Soranzo, N. The Impact of Rare and Low-Frequency Genetic Variants in Common Disease. Genome Biology. 2017. https://doi.org/10.1186/s13059-017-1212-4.

(21) Mousavizadeh, L.; Ghasemi, S. Genotype and Phenotype of COVID-19: Their Roles in Pathogenesis. Journal of Microbiology, Immunology and Infection. 2020. https://doi.org/10.1016/j.jmii.2020.03.022.

